# Calicivirus VP2 forms a portal to mediate endosome escape

**DOI:** 10.1101/397901

**Authors:** Michaela Conley, Marion McElwee, Liyana Azmi, Mads Gabrielsen, Olwyn Byron, Ian G. Goodfellow, David Bhella

## Abstract

To initiate the infectious process, many viruses enter their host cells by triggering endocytosis following receptor engagement. The mechanism by which non-enveloped viruses, such as the caliciviruses, escape the endosome is however poorly understood. The *Caliciviridae* include many important human and animal pathogens, most notably norovirus, the cause of winter vomiting disease. Here we show that VP2, a minor capsid protein encoded by all caliciviruses, forms a large portal assembly at a unique three-fold symmetry axis following receptor engagement. This feature surrounds an open pore in the capsid shell. We hypothesise that the VP2 portal complex is the means by which the virus escapes the endosome, penetrating the endosomal membrane to release the viral genome into the cytoplasm. Cryogenic electron microscopy (cryoEM) and asymmetric reconstruction were used to investigate structural changes in the capsid of feline calicivirus (FCV) that occur when the virus binds to its cellular receptor junctional adhesion molecule-A (fJAM-A). Near atomic-resolution structures were calculated for the native virion alone and decorated with soluble receptor fragments. We present atomic models of the major capsid protein VP1 in the presence and absence of fJAM-A, revealing the contact interface and conformational changes brought about by the interaction. Furthermore, we have calculated an atomic model of the portal protein VP2 and revealed the structural changes in VP1 that lead to pore formation. While VP2 was known to be critical for the production of infectious virus, its function has been hitherto undetermined. Our finding that VP2 assembles a portal that is likely responsible for endosome escape represents a major step forward in our understanding of both the *Caliciviridae* and icosahedral RNA containing viruses in general.

## Introduction

Many viruses enter their host cells by triggering endocytosis following receptor engagement [1]. Enveloped viruses, such as influenza A virus, escape the endosome by fusing their viral membrane with the endosomal membrane [2]. This process is mediated by dedicated fusion proteins embedded in the viral envelope and leads to the formation of a pore through which the virion contents pass into the cytosol [3]. Despite the critical nature of this stage of the infectious process, little is known about mechanisms of endosome escape in non-enveloped viruses such as the caliciviruses.

The *Caliciviridae* family of viruses comprises five genera: *Vesivirus, Norovirus, Sapovirus, Lagovirus* and *Nebovirus*. Noroviruses and sapoviruses cause outbreaks of acute gastroenteritis in humans, known as winter vomiting disease [4, 5]. Vesiviruses include several important veterinary pathogens such as feline calicivirus (FCV) a major cause of respiratory disease in felids [6]. Caliciviruses have positive sense RNA genomes of approximately 7.5 kilobases that are packaged within a spherical capsid of ∼40nm diameter [7]. Their genomes contain between two and four open reading frames (ORFs). ORF1 encodes the non-structural viral proteins, expressed as a polyprotein that is posttranslationally cleaved to yield the individual proteins. In noroviruses and vesiviruses ORF2 encodes the major capsid protein VP1 which, in other caliciviruses, is encoded in ORF1. The second/third ORF encodes the minor capsid protein, VP2 [8]. Murine norovirus is known to contain a fourth ORF that encodes the VF1 protein involved in evasion of innate immunity [9].

Calicivirus capsids have a characteristic virion morphology; the T=3 icosahedrally symmetric shell assembles from 180 copies of VP1 in three quasi-equivalent settings. Termed A, B and C, these VP1 conformers form 90 dimeric capsomeres. AB capsomeres are arranged around the 5-fold symmetry axes while CC capsomeres are located at 2-fold symmetry axes, resulting in alternating AB and CC dimers about the 3-fold symmetry axes [7, 10]. The VP1 protein of vesiviruses is proteolytically processed after translation, removing the N-terminal 125 residues. The mature capsid protein has a short N-terminal arm (NTA), a shell (S) domain comprising an eight-stranded antiparallel β-barrel motif, which forms the floor of the capsid, and a protruding (P) domain. The P domain dimers are responsible for the characteristic calicivirus morphology, presenting as broad arch-shaped spikes that, at low-resolution give the impression of 32 cup-shaped depressions on the virus surface (hence *calici*-from the greek *calyx* or cup). The P domain is divided into P1 and P2 sub-domains with P2 occurring as an insertion in P1 and forming the outer surface of the capsid protrusions. In addition to VP1, a hitherto unknown but low number of VP2 molecules are incorporated into the virion [11]. VP2 has been shown to be essential for the production of infectious virions [12], and it has been suggested that the small (12kDa) protein may play a critical role in the assembly of progeny virus particles [13]. It has been hypothesised to bind to viral RNA owing to the presence of a large number of basic amino-acids [8]. Norwalk virus VP2 has been shown to interact with VP1. An isoleucine residue at position 52 was found to be essential for this interaction and has been mapped to the inner surface of the capsid [14]. Despite considerable effort, no calicivirus structures solved to date present density assigned as VP2.

Feline calicivirus represents an excellent model system for the study of calicivirus biology as it is readily propagated *in vitro* and is one of only three caliciviruses to date for which a protein receptor has been identified [15-17]. FCV binds to the tight-junction cell-adhesion molecule feline junctional adhesion molecule-A (fJAM-A). JAM-A is composed of a short cytoplasmic tail, a transmembrane domain and two immunoglobulin-like domains; the membrane distal V-type D1 and proximal I-type D2. Although no structure for fJAM-A is available, X-ray crystallography studies of the murine and human homologues have shown them to form U-shaped dimers via *cis*-interactions at the GFCC’ face of D1 [18, 19]. JAM-A is also predicted to form interactions in *trans* across tight-junctions. We have previously shown that the outer face of the FCV P2-domain binds to D1 of fJAM-A in a monomeric state. The site of FCV binding is however on the other side of fJAM-A D1, away from the homo-dimerisation site [20, 21]. Our analysis revealed that upon receptor engagement, the FCV capsid undergoes considerable conformational changes; the capsid spikes rotate approximately 15° counter-clockwise. Based on these data, we hypothesised that the conformational change was a priming step that would prepare the virion for genome release following internalisation. FCV enters cells via clathrin mediated endocytosis. Endosomal acidification has been shown to be a necessary step in the viral entry process although the mechanism by which the virus escapes the endosome, delivering its genome into the cytoplasm has until now not been elucidated[22]. Interestingly there is a growing body of evidence to suggest that the extent of conformational changes subsequent to fJAM-A binding may correlate with pathogenicity in FCV [10, 23].

Here we present high-resolution cryoEM structures for FCV both undecorated and labelled with a soluble ectodomain fragment of fJAM-A. We show that upon receptor engagement significant conformational changes occur in the virion, leading to the formation of a large funnel-shaped portal structure at a single unique three-fold symmetry axis. The portal, which was not detected in undecorated virions, is formed of twelve copies of the minor-capsid protein VP2 arranged with their hydrophobic N-termini pointing away from the virion surface. Local rearrangement of VP1 at the portal site leads to opening of a pore in the capsid shell. We hypothesise that the portal assembly functions as a channel for delivery of the calicivirus genome through the endosomal membrane into the cytoplasm of a host cell to initiate infection.

Our study presents the first description of a portal assembly in a positive-sense RNA-containing animal virus and provides insights into a critical step in the replication cycle of an important group of viral pathogens, that may serve as a paradigm for endosome escape in other RNA containing icosahedral viruses.

## Results

### Cryo-electron microscopy of feline calicivirus decorated with a soluble fragment of feline junctional adhesion molecule-A

To construct an atomic resolution model of FCV interacting with its cellular receptor fJAM-A and resolve the molecular detail of conformation changes induced, we collected high-resolution cryo-EM images of FCV both decorated with fJAM-A and undecorated. 5,198 micrographs of FCV and 13,865 micrographs of FCV-fJAM-A were recorded on a Thermo-Fisher Titan Krios microscope equipped with a Falcon III camera (figure 1a,b). 41,436 particle images of undecorated FCV virions were processed to calculate an icosahedrally symmetric reconstruction having a nominal resolution of 3 Å (figures 1c,d, S1, movie S1). 71,671 particle images of fJAM-A decorated FCV virions were likewise processed to calculate an icosahedral reconstruction of the virus-receptor complex, achieving an overall resolution of 3.5 Å (figure 1e,f, S2, movie S1). Inspection of sections through the reconstructed density map, and local-resolution analysis confirm our previous findings that conformational changes following receptor engagement result in blurring of features in the P-domain and receptor density, preventing us from achieving a high-resolution view of this region. To address this, we have previously performed model-based classification of the conformational changes induced in the virion following receptor engagement [21]. Using the resulting density maps of FCV-fJAM-A in pre and post-conformational change states we repeated this analysis on our high-resolution dataset. While we were able to extract a very small sub-set of particles (493 out of 71,671) that were found to be in a pre-conformational change state, i.e. all of the capsomeres were in the same orientation as the undecorated virus (figure S3), the vast majority of the dataset were grouped into classes in which the P-domains remained poorly resolved. Thus, we conclude from this analysis that the conformational change is not coordinated. This view is supported by our finding that at the resolution of this study, there are no appreciable differences in the structure of the S-domain of VP1 in our icosahedral reconstructions of FCV in the presence or absence of fJAM-A.

**Figure 1.**
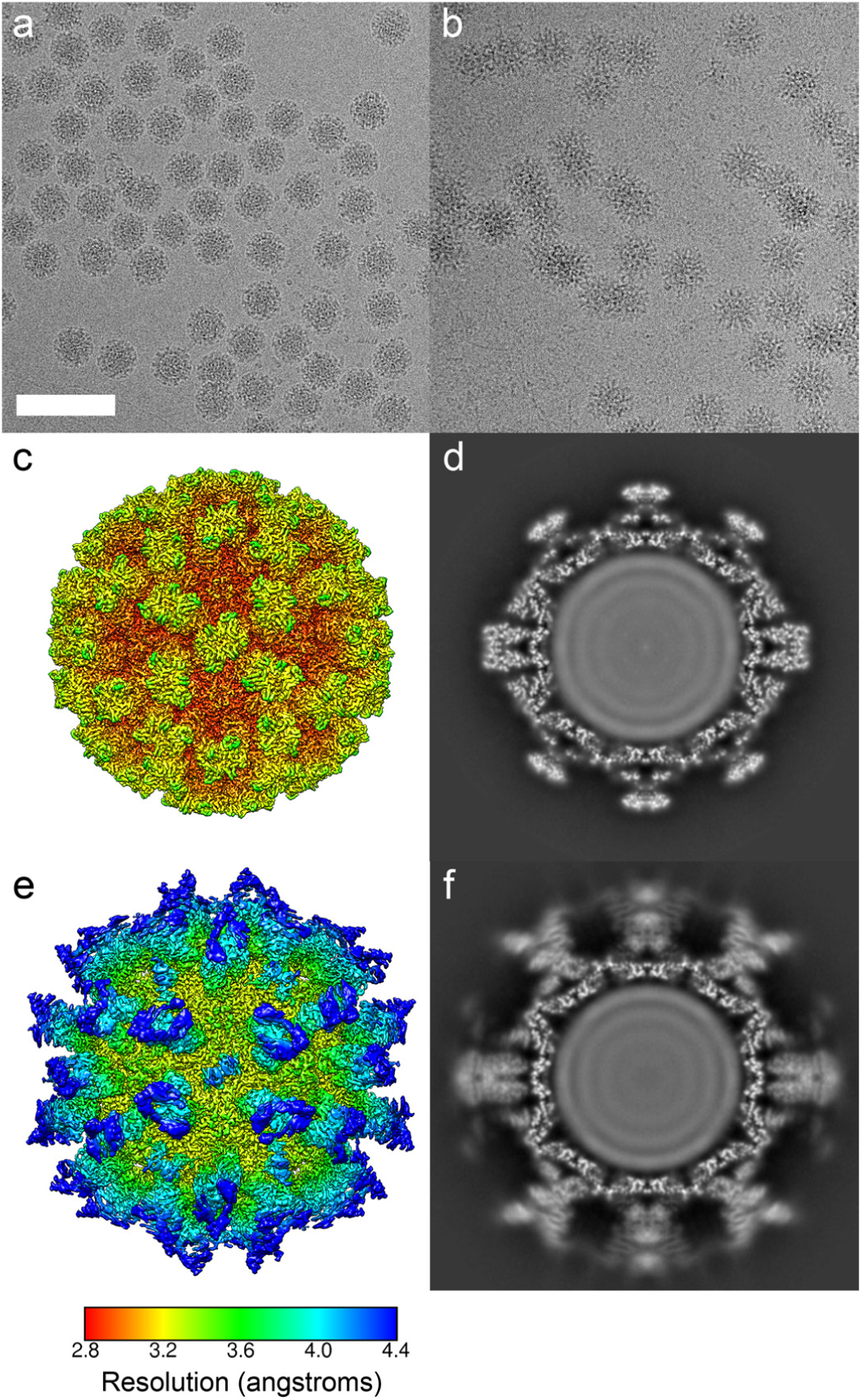
Cryogenic electron microscopy of feline calicivirus (FCV) strain F9 virions both unlabelled (a) and decorated with soluble ectodomain fragments of feline-junctional adhesion molecule A (fJAM-A) (b), scale bar is 100 nm. An icosahedral three-dimensional reconstruction of the unlabelled virion structure was calculated and achieved an overall resolution of 3 Å. An isosurface thresholded view is shown coloured and filtered according to local resolution, showing that in the shell of the capsid the resolution approaches 2.8 Å, while on the outer faces of the protruding capsomeres the resolution is closer to 3.5 Å (c). A central section through the reconstructed density map shows that both the shell (S) and protruding spike (P) domains are sharply resolved. Icosahedral reconstruction of receptor decorated FCV virions resulted in a lower overall resolution of 3.5 Å. The isosurface representation (e) is similarly coloured and filtered according to local resolution, revealing that in this map the dimeric P-domain spikes and fJAM-A fragments are resolved at poorer than 4 Å resolution. A central slice through the reconstructed density map shows that while the capsid shell is sharply resolved, the P-domains and fJAM-A components are blurred as a consequence of the receptor induced conformational changes in this region (f).

### Asymmetric reconstruction of AB and CC capsomeres

To calculate 3D reconstructions at sufficiently high resolution to allow building of atomic models of FCV P-dimers bound to fJAM-A in multiple conformations, we implemented a focussed classification method. Rather than calculating reconstructions of the whole virion, we set out to reconstruct individual capsomeres as they undergo receptor induced conformational changes, breaking icosahedral symmetry. Focussing the analysis on single AB or CC dimer positions in the capsid, we assigned data to self-similar classes for 3D reconstruction. This approach yielded 3D reconstructions with improved resolution and revealed the range and extent of conformational changes (figures S4-5, movie S2). Both AB and CC dimers were seen to present a variety of conformations upon fJAM-A engagement. The best resolved were those in which the P-domains were rotated and tilted towards the three-fold symmetry axes. Coherent averaging of structurally similar or identical capsomeres yielded sharper density maps. In some classes the P-fJAM-A density was resolved at resolutions of between 4 and 5 Å and was deemed suitable to be interpreted for the purposes of model building.

### A large portal assembly at a unique three-fold symmetry axis

In the course of the focussed classification analysis, three intriguing classes emerged in which we saw a previously unreported structure. Twelve fingers of density were seen to protrude from the capsid floor, arranged about the three-fold symmetry axis (figures S4-5, movie S2). Furthermore, the P-fJAM-A density in these classes was determined at improved resolution, suggesting that the capsomeres were constrained by the presence of the novel feature. To see this feature more clearly, focussed classification was performed on the three-fold axes. This revealed the presence of a large dodecameric portal-assembly extending approximately 13 nm from the capsid shell (figure 2, S6, movie S3). The portal protein monomers are arranged in a circle about the icosahedral three-fold symmetry axis, forming a funnel-shaped structure that is 7nm in diameter at its base and 5nm in diameter at its tip, where the density becomes fuzzy and poorly resolved. The portal protein density indicates a predominantly α-helical structure and is well resolved proximal to the capsid shell (figure 2d). Interestingly, there is also an opening in the capsid shell at the portal associated three-fold symmetry axis that is not seen in non-portal axes (figure 2c, S6d and movie S3).

**Figure 2.**
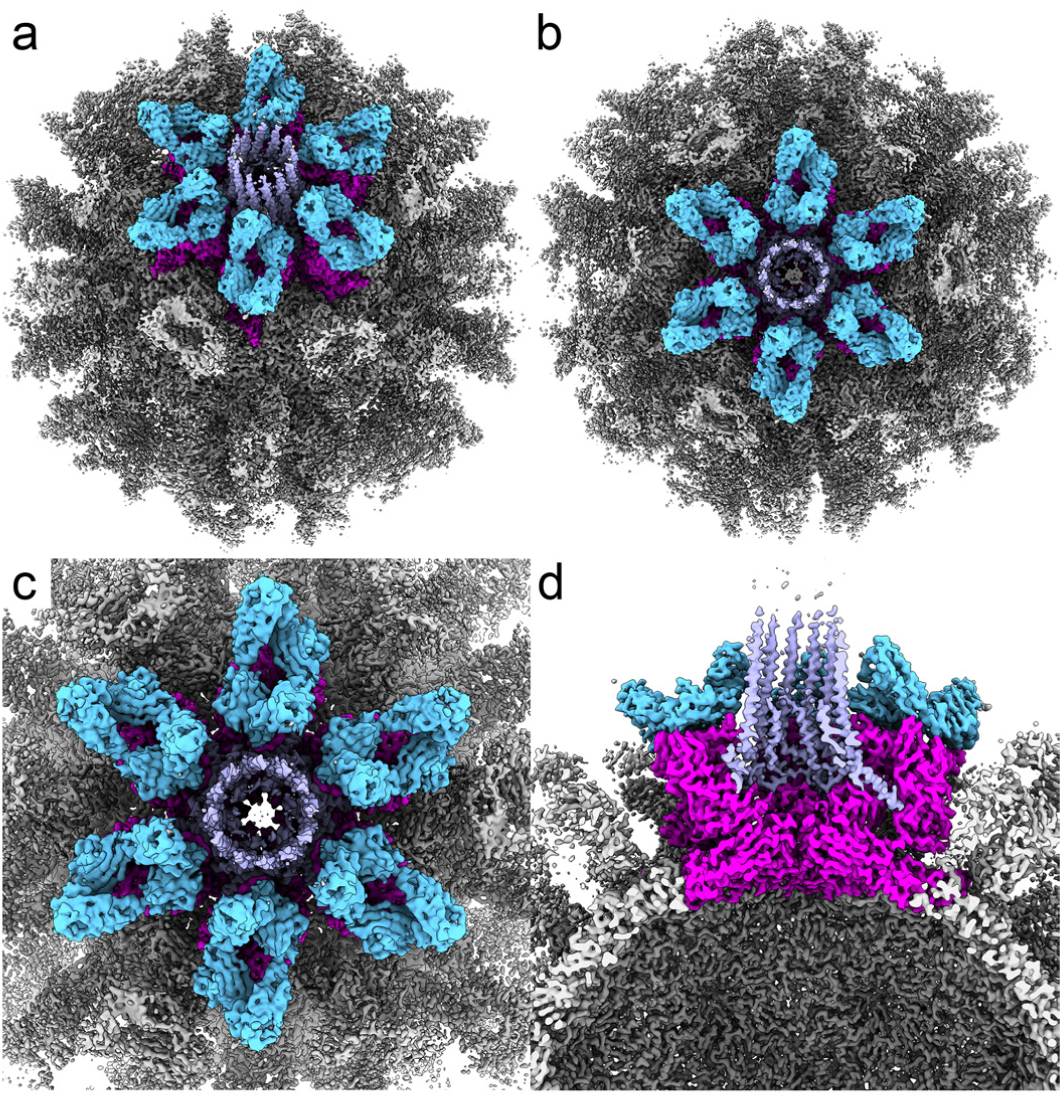
C3-symmetric 3D reconstruction of FCV decorated with fJAM-A following focussed classification analysis reveals the structure of a portal assembly at a unique three-fold symmetry axis. Ambient occlusion rendering of the isosurface thresholded reconstruction coloured to highlight the portal vertex viewed along the two-fold (a) and three-fold (b) symmetry axes. A close-up view of the portal assembly highlights the presence of a pore in the capsid shell at the centre of this symmetry axis (c). Cut-away view perpendicular to the portal axis (d). The map is coloured to highlight individual components: pink – VP1, blue – fJAM-A and lilac – VP2.

To calculate this reconstruction, we applied a focussed clas-sification approach to identify and reconstruct the unique three-fold portal-vertex in every particle image. Each of the 60 possible orientations for every particle (corresponding to the 60-fold redundancy of an icosahedral object) was subjected to 3D classification, grouping the data into self-similar classes based on features present at a single three-fold axis. Our data set of 71,671 particle images therefore gave rise to 4,300,260 views that were tested. Of these, 234,076 particle views were assigned to the class in which the portal assembly was present, corresponding to 5.44% of the dataset. This is consistent with each particle having a single portal at a unique three-fold axis, as there are 20 three-fold symmetry axes per virion. To further evaluate the likely number of portals per virion, we interrogated the metadata file that contained particle orientations for this class, to determine how many times each particle contributed to the 3D reconstruction (figure S7). This revealed that the median number of views contributed by each particle was 3; corresponding to the C3 symmetry of the object. Our data indicate that in the majority of particles a single portal is present at a unique three-fold axis.

Having established the presence of a novel structure on the surface of FCV decorated with soluble fJAM-A, we sought to determine whether it was present in undecorated particles. Having performed the same focussed-classification analysis on our undecorated dataset however, no such feature was found. This indicates that the portal assembly most likely emerges from within the virion following receptor engagement, possibly passing through the pore formed in the capsid shell.

Owing to the size of the portal proteins, we postulated that they were the minor capsid protein VP2, rather than a rearrangement of the major capsid protein VP1. As the portal and surrounding capsid and receptor densities were resolved at between 3 and 5 Å, we proceeded to build an atomic-resolution model of the assembly.

### An atomic model of the FCV strain F9 major capsid protein VP1 prior to receptor engagement

A structure of the FCV virion has been previously determined by X-ray crystallography [10]. In that study however, the analysis was performed on the virulent systemic (VS) FCV strain 5, rather than the vaccine strain F9 used in the current study. Differences between the sequences for VP1 in these two strains led us to build a new model based on our cryoEM density map. To achieve this, a previously calculated homology model of the FCV F9 VP1 major capsid protein was docked into our cryoEM map of the undecorated virus [20]. The model was manually edited to improve the fit to our density map and rectify sequence differences between our virus and the published sequence we used previously (figure S8). The model was then iteratively refined (table S1). The structure closely matched that of the previously solved FCV capsid protein, with the most significant differences being in the P2 domain [10]. Our model shows the canonical eight-stranded anti-parallel β-barrel, or ‘β-jellyroll’ motif in the S-domain (figure 3, movie S4). This is found in many viral capsid proteins and in particular, those of positive-sense RNA viruses. The P2 domain also incorporates an anti-parallel β-barrel that has been previously described in *Caliciviridae* capsid structures. The motif comprises six-strands and is cylindrical in shape. Interestingly within this feature we see density consistent with the presence of a metal ion, something that has not been reported in any calicivirus capsid proteins to date (figure 3b,c, movie S4). The majority of coordinating interactions occur with main-chain carbonyl groups, but side-chains of Q474 and D479 also play a role. The distances between the putative metal-ion density and surrounding oxygens (∼2.8 Å), along with the preponderance of main-chain carbonyl interactions, lead us to suggest that potassium is the most likely candidate for these densities [24]. At this resolution, however, it is not possible to unambiguously position all relevant atoms. A recent mutational analysis of FCV VP1 fJAM-A binding identified several amino-acid residues within or close to the coordinating sphere of this metal-ion, that were critical to virus infectivity. Viruses in which these sites were mutated (I482, K480 and H516) were able to both assemble capsids and bind to fJAM-A, but were not infectious [23]. Sequence analysis also highlighted differences between VS-FCV and non-VS strains within this region, including residue D479, which in VS-FCV strain 5 is an asparagine residue and is oriented away from the site of metal binding we see in strain F9 [10] (figure S9). These data suggest that the metalbinding site may be important for infectivity and pathogenesis.

**Figure 3.**
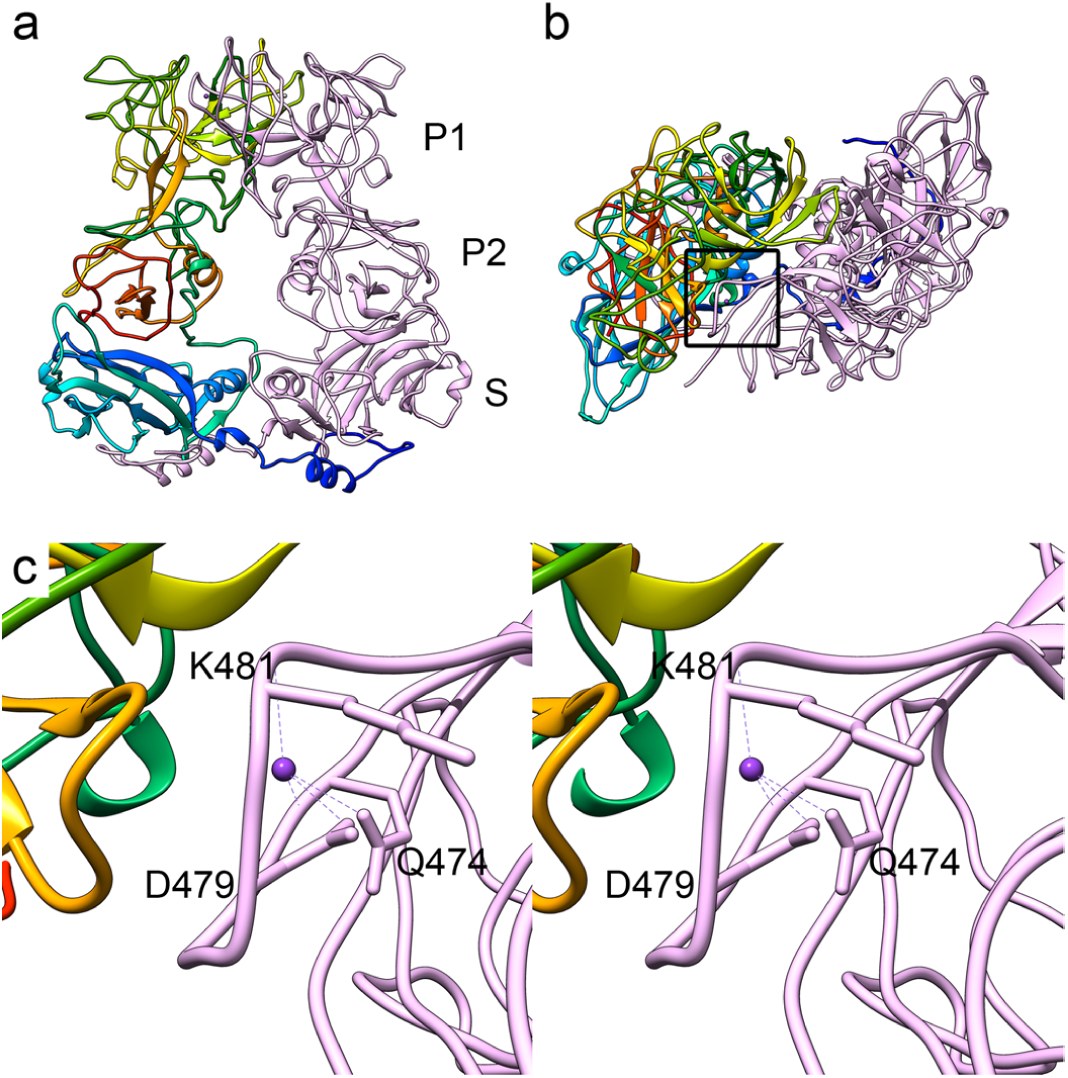
Ribbon diagrams to show the atomic model of the FCV strain F9 major capsid protein VP1, calculated by modelling the protein sequence into the icosahedral reconstruction of the undecorated virion. Chain A is coloured pale pink, chain B is presented in rainbow representation (N-terminus blue, C-terminus red). A side-view of the dimeric capsomere is labelled to identify the shell (S) and protruding (P1 and P2) domains (a). The S domain shows the characteristic ‘β-jellyroll’ motif seen in all calicivirus structures solved to date. A top-view of the VP1 dimer is shown with a box to highlight the location of a putative metal ion (b). A close-up, wall-eyed stereo view of this region is presented identifying the coordinating interactions within the metal binding site. Based on the distances measured for these interactions we suggest that this metal ion may be potassium (c).

### An atomic-resolution model of fJAM-A bound to VP1

To build a model for the structure of FCV following receptor engagement, the coordinates for FCV VP1 in the absence of fJAMA were divided into individual P and S domains for each of the three quasi-equivalent positions in the T=3 virus (A, B and C). These were docked to the appropriate regions of density in our 3D reconstruction of the unique portal-vertex of FCV-fJAM-A, to produce an asymmetric unit comprising one AB-dimer and one CC-dimer. To distinguish chains in the CC dimer, we designated the VP1 protomer proximal to the portal axis as chain D. Four copies of our previously calculated homology model for fJAM-A were likewise docked to our density map, as individual domains (D1 and D2). Both VP1 and fJAM-A models were manually edited and then refined to produce the final set of coordinates for the FCV-fJAM-A complex (table S2). These data provide a detailed view of the virus-receptor interface and structural changes in VP1 that occur following receptor engagement, leading to the formation of a pore in the capsid shell.

### fJAM-A VP1 interactions

The interactions between fJAM-A and VP1 were characterised for each of the four VP1 chains in the asymmetric unit, using tools in UCSF Chimera[25] and the ‘Protein interfaces, surfaces and assemblies’ service PISA at the European Bioinformatics Institute (http://www.ebi.ac.uk/pdbe/prot_int/pistart.html) [26] (figure S10, tables S3a-h, movie S5). The footprint of each fJAMA molecule is between 984 Å^2^ (fJAM-A^D^ – the molecule bound primarily to chain D) and 1068 Å^2^ (fJAM-A^A^). Interactions at the interface between virus and receptor are largely electrostatic in character. Between 1 and 6 hydrogen bonds are predicted to form between the capsid and each receptor fragment, these include one or more bonds formed between residues N444 and D445 in VP1 and Y51 and F54 in fJAM-A. These interactions are of particular interest as a loop on the outer face of VP1 at residues 436-448 undergoes significant rearrangements upon fJAM-A binding. It is notable that residues K480 and K481 are also identified as interacting with fJAM-A. K481 is highlighted as potentially forming a salt-bridge with E34 and a hydrogen bond with S33 of fJAM-A. These lysine residues lie within the putative metal binding site of VP1.

### Structural rearrangements in VP1 leading to portal assembly

We have previously shown that upon receptor engagement FCV undergoes conformational changes that we had proposed to be important for virus entry [20, 21]. In this study, we set out to produce an atomic level description of these structural changes. In doing so, we discovered that in addition to the previously identified capsomere rotations, receptor binding leads to the formation of a pore in the capsid shell at a unique three-fold axis, and assembly of a large portal comprising twelve copies of the minor capsid protein VP2. To provide a detailed understanding of the structural changes in VP1 wrought by fJAM-A binding we aligned the P domains of undecorated and fJAM-A labelled structures for both AB and CC dimers. This allowed us to focus our view on the subtler structural rearrangements of the P-domain, ignoring the large rotations relative to the S-domain (figure 4). Molecular dynamics was used to morph the aligned structures, further highlighting the nature of structural rearrangements that extend from the outer face of P2, through the P-domain to the hinge that connects the protruding capsomere to the capsid shell (movie S6). The most striking structural changes take place in the P2 domain and in particular in the abovementioned 436-448 loops, which rise 3 Å towards the receptor, to form an interaction that may involve several H-bonds.

**Figure 4.**
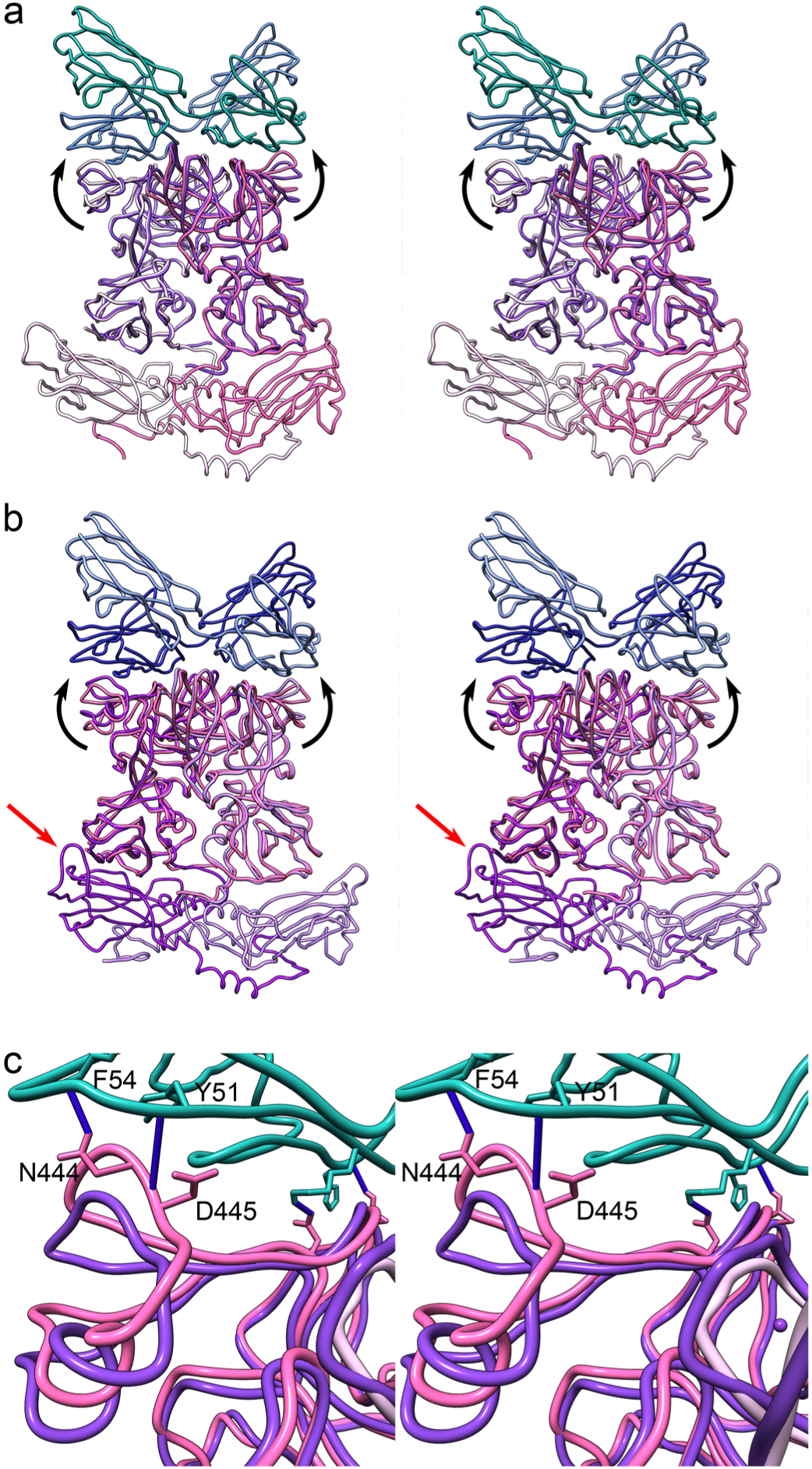
Ribbon diagrams to show the atomic models for FCV VP1 and fJAM-A following receptor engagement. Wall-eyed stereo views are shown for the AB-dimer (a) and CC-dimer (b). In panel (a), chain A is coloured pale pink, while chain B is coloured hot pink, fJAM-A^A^ is coloured blue, fJAM-A^B^ is coloured green. Overlaid on the structure is the aligned AB P-domains of the undecorated virus (purple), highlighting the structural rearrangements that occur following fJAM-A binding. The most striking structural change is the upwards movement of loop 436-448, indicated by black arrows. Panel (b) shows the structure for the CC dimer, the portal proximal chain (now designated chain D) is coloured dark purple, while the distal chain C is coloured pale purple, and fJAM-A molecules are coloured blue. As in panel (a), the structure of VP1 in the absence of receptor is overlaid (pink) to highlight the structural rearrangements that occur following receptor engagement. This representation of the CC dimer also highlights the structural rearrangements that occur in the S-domain leading to the opening of a pore in the capsid shell. Compare the orientation of the 293-307 loop in chain D (red arrow) to the same loop in chain C. (c) A close-up view of the 436-448 loop of chain B is shown highlighting the H-bonds that are predicted to form between VP1 and fJAM-A (dark blue).

Inspection of the structures of VP1 before and after receptor engagement also reveals the structural rearrangements at the icosahedral three-fold axis that lead to the opening of a pore in the capsid shell (figure 4b, 5). Consistent with the T=3 structure of the FCV capsid, the three-fold axes exhibit quasi-six-fold symmetry. Chains B and C alternate about this axis and six tyrosine residues (Y300) from these chains point inwards, with the side chains of Y300_B_ tilted towards the capsid interior and those of Y300_C_ tilted towards the exterior. Upon receptor engagement the loops bearing these residues (293-307) fold outwards, resulting in the opening of a pore in the capsid shell that is ∼1 nm in diameter (movie S7). A solvent-excluded surface representation of the atomic model also shows that the counter-clockwise rotation of the capsomeres arranged about the portal vertex, following receptor binding, leads to a closing together of the P1 domains, forming a closed cup around the pore (figure 5b).

**Figure 5.**
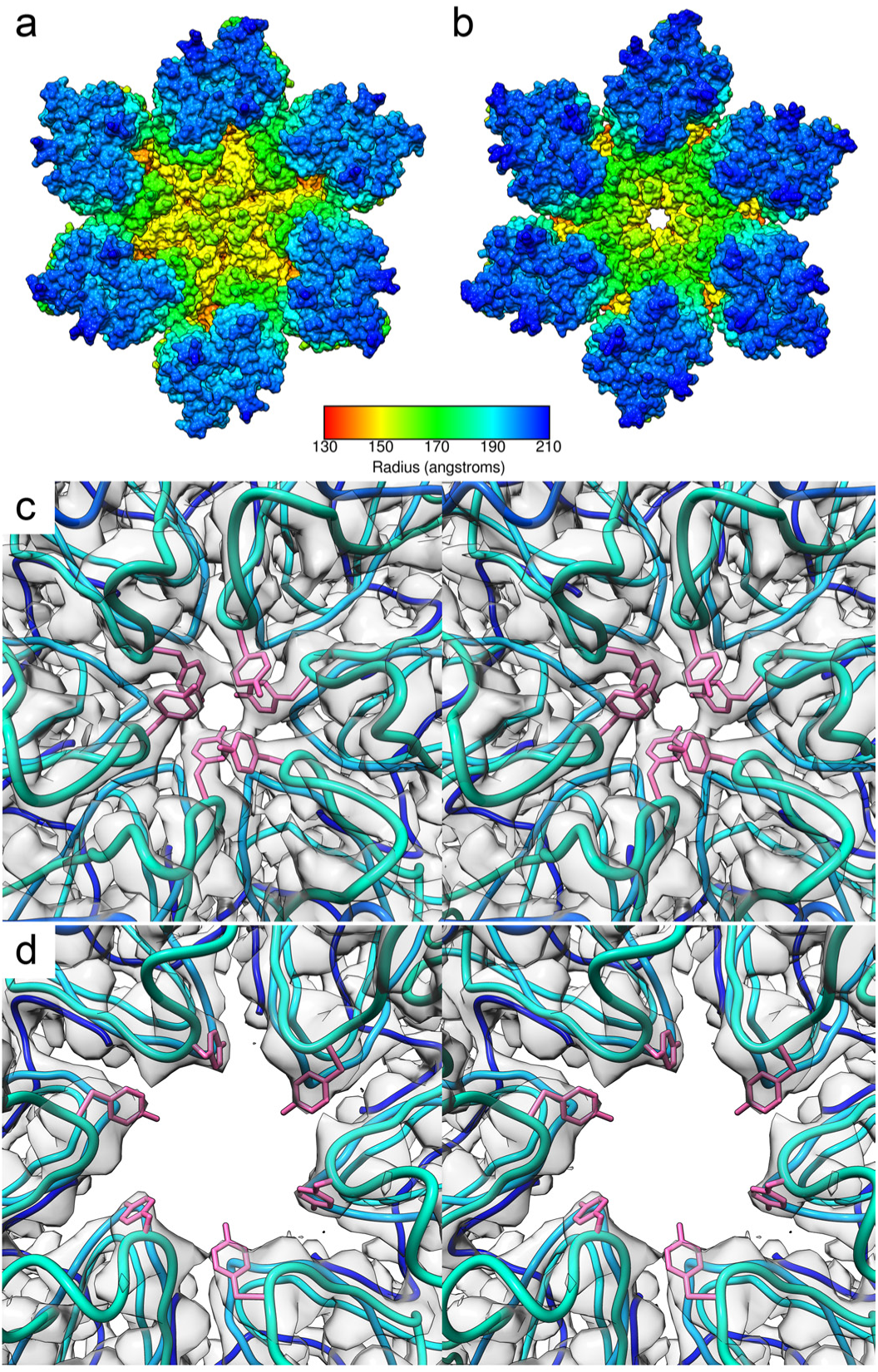
Conformational changes in VP1 following receptor engagement leading to the formation of a pore in the capsid shell. Solvent-excluded surfaces of VP1 dimers arranged about the three-fold symmetry axis in undecorated FCV (a), and fJAM-A labelled FCV at the portal axis (b) – coloured by radius. A close-up, wall-eyed stereo view of the icosahedral three-fold axis in the undecorated FCV reconstruction viewed from the capsid exterior (c) shows that this axis is formed of six tyrosine (Y300) side chains that point to the centre of the symmetry axis. Three tyrosine residues are each donated by chains B and C, that alternate about the symmetry axis following the quasi-six-fold symmetry seen in a T=3 icosahedral structure. Y300_B_ points towards the capsid interior, while Y300_C_ points outwards from the capsid surface. Following receptor engagement, a portal complex assembles at a unique three-fold axis. At the centre of this axis, a pore opens in the capsid surface by the folding outwards of loop 293-307 (d). Panels (c) and (d) present both modelled coordinates (ribbon diagram in rainbow colour scheme, atomic representation in pink) and the reconstructed density map (transparent grey).

### The structure and assembly of the calicivirus portal protein VP2

Following modelling of VP1 and fJAM-A, the amino-acid sequence of VP2 was built ab initio into the remaining density. Many bulky side-chains were clearly resolved and allowed us to construct a robust model, particularly in the regions proximal to the capsid (figure 6 and movie S8). This confirmed our hypothesis that the portal assembly consists of the minor capsid protein VP2. Twelve copies are arranged about the unique three-fold axis to form a funnel shaped tube. The asymmetric unit therefore includes four copies of VP2 (designated as chains I-L). The structure of VP2 includes three α-helices. The first and longest α-helix (*a*) makes up the body of the funnel and extends from residue 18 to 62. The N-terminal 17 residues are not resolved and are distal to the capsid surface. Interestingly these N-terminal residues of VP2 are characterised as being highly hydrophobic by Kyte-Doolittle analysis (figure S11). VP2 is present in two distinct conformations that present different folds for residues 75-106. The two conformers alternate about the portal axis (six copies of each conformer). In conformer one (chains J and L), α-helices *b* (61-74) and *c* (91-106) bind to the capsid P domains via electrostatic interactions (table S4a-h). The C-terminal α-helices (*c*) have positively charged surfaces that pack against the P1-domains of both VP1 protomers in the dimeric capsomeres that alternate about the portal axis (AB or CD), binding within a negatively charged cleft to anchor the portal assembly to the capsid. The *bc* loops (residues 75-90) wrap across the surface of the adjacent VP1 molecule laying closest to the portal axis (chains D or B respectively), and helix *b* binds into a second negatively charged cleft on the surface of the P2 domain that is formed by the rearrangements of loop 436-448 subsequent to fJAM-A binding. This interaction is predicted to include multiple salt bridges and hydrogen bonds (figure 6, tables S4a-h, movie 8). Thus, each of the VP2 molecules in the first conformation binds to three VP1 molecules: VP2_J_ is anchored to VP1_AB_ in the P1 binding site. The *bc* loop then wraps across the surface of VP1_D_ and helix *b* inserts into the second binding site, in the P2 domain of VP1_D_. Likewise, helix *c* of VP2_L_ binds to VP1_CD_ in the P1 cleft. It then folds across the face of VP1_B_ binding into the second interaction site on that protomer. In the second conformation of VP2 (designated as chains I and K) the C-terminal residues (75-106) do not bind VP1, rather they fold into the lumen of the portal, such that six symmetry related C-terminal α-helices form an inner ring of fingers pointing away from the capsid surface. The C-terminal 6 residues of this conformer are not resolved. Conformer two barely contacts VP1 at all, the contact interface between this conformation and the capsid is ∼150Å^2^. VP2 is instead held within the portal complex primarily by hydrophobic interactions along the interface between the N-terminal α-helices (*a*). Interestingly, the C-terminal region of this conformer presents a largely negatively charged surface to the interior of the portal.

**Figure 6.**
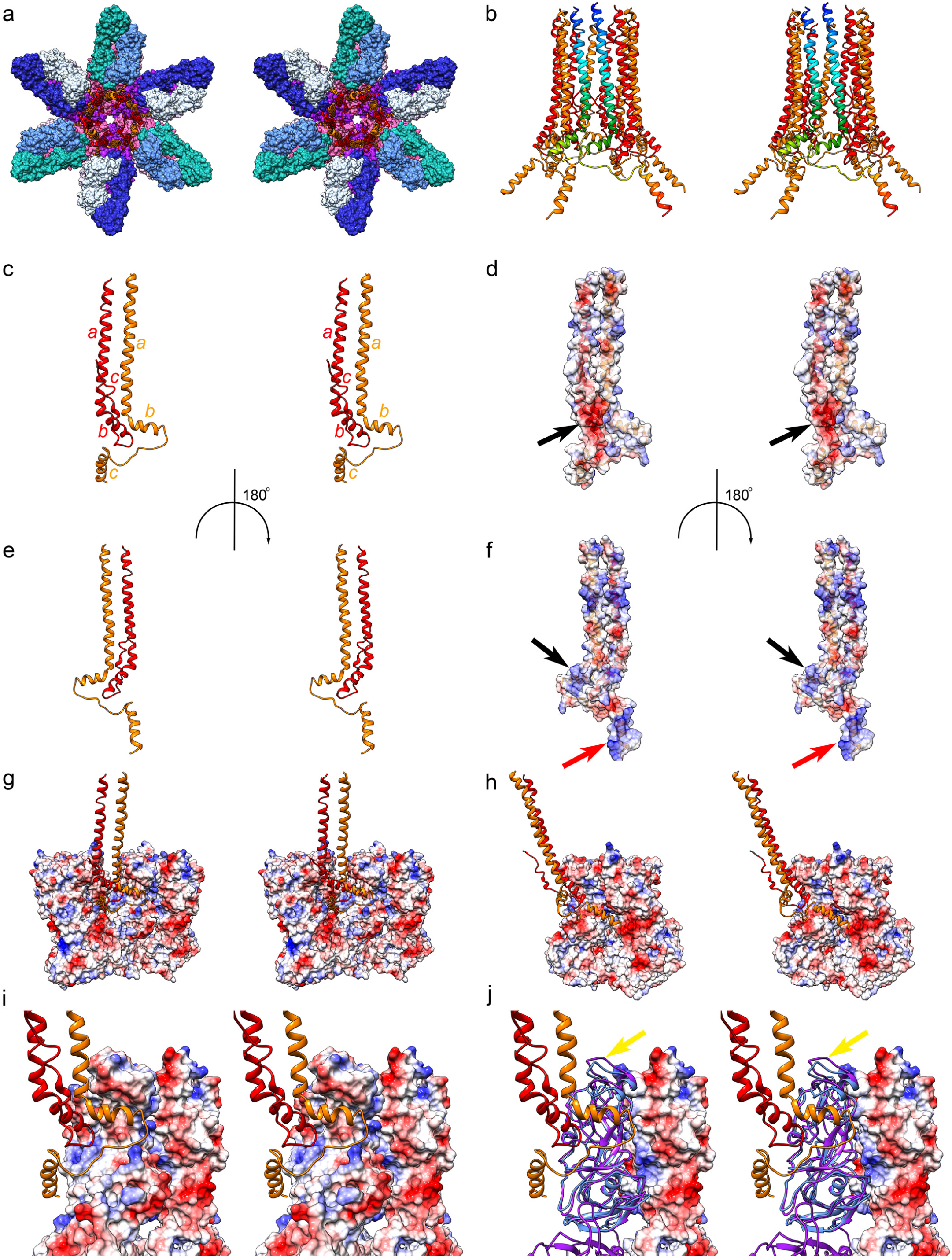
Wall-eyed stereo pair images of VP2 to show the folds of the two conformers that are seen to alternate about the three-fold portal axis and highlight the interactions between conformer one (orange) and the major capsid protein VP1. A solvent-excluded surface representation of the portal vertex is presented viewed along the unique three-fold axis. Six dimers of VP1 are shown, AB (pink) and CD dimers (purple) alternate about the symmetry axis. On the outer surface of the P2 domain, fJAM-A molecules are bound (blue/green), with one receptor molecule binding to each VP1. The portal assembly, comprising twelve copies of the VP2 protein is shown as a ribbon diagram (orange/red). The VP2 dodecameric portal structure is shown alone, viewed perpendicular to the portal axis (b). VP2 has three α-helices designated *a-c*. Two conformers of the molecule are seen in this assembly, conformer one (orange) and conformer two (red). The front most VP2 dimer is shown in rainbow colouring, highlighting that the N-terminus of VP2 is distal to the capsid shell. The C-terminal residues (75-106) present different folds in the two conformers, with conformer one showing a well-ordered helix *c* that splays outwards, while the C-terminus of conformer two folds into the lumen on the portal. A dimer of VP2 showing the two conformations is presented viewed from the portal interior, α-helices are labelled. The poorly-ordered C-terminal helix *c* of conformer two can be seen leaning towards the viewer (c). A solvent-excluded surface representation of this view, coloured to show surface potential (negative = red, positive = blue), shows that the C-terminal region of conformer two presents a negatively charged surface to the portal interior (black arrow – d). Panels e and f show the same VP2 dimer rotated 180° about the vertical axis, the outward facing C-terminal helix of conformer one is now leaning towards the viewer. Interestingly, both helix *b* and *c* in this conformer present positively charged surfaces (black and red arrows respectively), which bind to negatively charged clefts on the capsid surface (g). Two binding sites are present on the capsid surface, helix *c* binds to the P1 domains of both VP1 molecules in each dimeric capsomere arranged about the portal axis (h), while helix *b* binds to a cleft on the adjacent portal proximal VP1 molecule in the P2 domain (i). This binding site is opened up by the upwards movement of loop 436-448 following fJAM-A binding. Ribbon diagrams of the portal proximal VP1 molecule (in this case chain D) are shown for both fJAM-A decorated (purple) and unlabelled (blue) VP1, showing that without the structural rearrangements of loop 436-448 brought about by receptor binding, the helix and loop (450-460) laying immediately below clashes with helix *b* of VP2 (yellow arrow) (j).

### fJAM-A binding renders FCV virions unstable at low pH

The FCV entry pathway has been shown to utilise clathrin-mediated endocytosis[22]. We therefore set out to test whether RNA release could be triggered by reduction in pH, and whether this would be fJAM-A dependent. An RNA release assay was devised in which virus was incubated in the presence of the nucleic acid binding dye Syto9 at differing pH and in the presence or absence of fJAM-A. In this assay, viral uncoating was detected by fluorescence induced when Syto9 binds to the released RNA. We found that in the absence of fJAM-A, no RNA was released when pH values were varied from 3 to 9 (the limits of our experiment). In the presence of fJAM-A however RNA release was detected at pH3 and 4 (figure S12). Negative stain transmission electron microscopy revealed that at low pH, in the presence of fJAM-A, FCV virions disassembled (figure S13).

Our structural analysis indicates that upon fJAM-A binding to FCV, VP2 emerges from the virion interior to form a portal. We hypothesise that this is the means by which viral RNA is released from the endosome. Moreover H+ concentration in the late endosome is thought to reach a pH of ∼5, with further acidification to ∼pH 4.5 being achieved in the lysosome [27]. Thus, we do not suggest that virion disassembly under very low pH conditions is likely to play a role in genome release in vivo. Nonetheless, we consider this assay to be a useful tool to establish the stoichiometry of receptor binding that is sufficient to induce structural changes in FCV that are associated with entry. We therefore measured RNA release for FCV virions in the presence of varying quantities of the fJAM-A ectodomain at low pH to estimate the ratio of fJAM-A to VP1 needed to destabilise the capsid. The key ratios were estimated at between 1:9 and 1:14 (figure S14). We confirmed our findings by using negative stain electron microscopy to visualise particle disassembly at low pH (figure S15). In the presence of fJAM-A, no intact particles were visible at fJAM-A:VP1 ratios of between 1:1 and 1:10. Interestingly, in these images we saw small balls of density that may be condensed viral RNA. When a fJAM-A:VP1 ratio of 1:11 was applied at pH3, broken and irregular particles were observed, suggesting that 1:10 is the minimum ratio capable of destabilising all of the capsids.

### FCV disrupts fJAM-A homodimers

Our analysis of the structure of FCV bound to fJAM-A indicates that a single receptor molecule binds to each VP1 monomer. In structural studies of the human and murine orthologs, JAM-A is found to form dimers. To determine whether fJAM-A ectodomain fragments were dimeric in solution we used small-angle X-ray scattering (SAXS) to calculate a low-resolution model of the protein solution structure. Ab initio model-building yielded a shape consistent with fJAM-A dimer formation (figure S16, movie S9). Docking of the atomic models for both human and murine JAM-A into the SAXS envelope for fJAM-A indicates that the FCV receptor likely forms similar dimers in solution. We have previously noted that the FCV binding site on fJAM-A is on the opposite face of the receptor than the dimerisation site found in human and murine JAM-A, but that the dimerization site was occluded by the D2 domain of symmetry related fJAM-A molecules when bound to FCV [28]. Our high-resolution model of FCV decorated with fJAM-A reveals a significant contact interface between the two receptor molecules when bound to the capsid (672 Å2). Thus, a second dimerisation interface arises on fJAM-A as a consequence of FCV binding, this time forming between D1 and D2, as opposed to the homodimerisation site that occurs between D1 domains (table 5a-d). It therefore seems reasonable to suggest that fJAM-A homodimers at the cell surface will be disrupted by binding of FCV.

## Discussion

Our study provides the first description of the structure and function of calicivirus VP2, a minor capsid protein present in all caliciviruses including the medically important norovirus and sapovirus families. VP2 has long been known to be essential for the formation of infectious virus[12]. We have now demonstrated that VP2 functions as a portal that most likely emerges from the capsid interior following receptor engagement. Structural rearrangements of VP1 at the portal site lead to the formation of a pore in the capsid shell. The distal tips of VP2 are hydrophobic in character, strongly suggesting that the purpose of the VP2 portal is to insert into a membrane. Taken together our findings strongly suggest that the function of VP2 is to mediate endosome escape, forming a channel in the endosomal membrane through which the viral genome is released into the cytoplasm.

VP2 has been postulated to be an RNA binding protein [8]. To act as an effective mediator of genome release, we consider that it is indeed likely to associate with one end of the viral genome. There is however no evidence of an interaction between VP2 and VPg, a protein found covalently linked at the 5’ end of the viral genome that functions both to prime RNA synthesis and initiate translation [29-31]. VP2 is however, known to bind to the capsid interior in noroviruses[14]. We propose that VP2 likely binds both the viral genome and the capsid interior at a single three-fold symmetry axis. Upon receptor engagement conformational changes at the capsid exterior trigger rearrangement of VP1 loop 293-307 at the three-fold axis, to form a pore through which VP2 can exit the virion and assemble the portal structure. Rotation of the P domains about the portal axis, leads to a closing together of the P1 domains while structural rearrangements in the P2 domain of VP1 at loop 436-448 stabilise the interaction with fJAM-A and open a cleft on the surface of P2 to which helix b of VP2 binds. Once the first conformer of VP2 is anchored to the capsid, addition of VP2 molecules in their second conformation leads to the assembly of the complete portal.

We hypothesise that the loss of capsid stability at low pH is a result of the structural changes induced by receptor binding. Whether this is a consequence of VP2 portal assembly, pore formation or capsomere rotation is unclear. Nonetheless, our data delineate the ratios of receptor to capsid protein necessary to induce destabilisation in solution, requiring only 18 receptor molecules per virion.

Using small angle X-ray scattering, we have shown that fJAM-A ectodomain fragments exist as dimers in solution, whereas our cryoEM data indicate that they bind to FCV as a monomer such that the D2 domains of symmetry related receptor molecules disrupt dimerisation. This is of interest because it has previously been shown that addition of adenovirus fibre-knob protein can trigger endocytosis by disruption of homodimers of the immunoglobulin-like tight-junction protein CAR (coxsackie-adenovirus receptor) [32]. Thus, we suggest that disruption of fJAM-A homodimers at the cell surface may be the mechanism by which FCV triggers endocytosis.

Our data lead us to propose a mechanistic model of FCV entry and endosome escape: FCV binding to fJAM-A at the cell surface leads to internalisation by endocytosis, triggered by fJAMA homodimer disruption. Enwrapping of the virion within the endosome leads to further binding of fJAM-A molecules, triggering conformational changes in the capsid surface that result in formation of the portal vertex. The hydrophobic N-termini of VP2 insert into the endosomal membrane forming a channel through which the genome may be released.

In our study viral genomic RNA remained associated with the capsid following receptor engagement until the pH was lowered to 4 which caused virion disassembly. It is known that acidification of the endosome is essential for FCV entry [22]. Late endosome acidity does not exceed pH 5 however [27]. We therefore think it unlikely that virion disassembly is the mechanism by which the genome is released from the endosome. It is also notable that the pore we have identified in the capsid shell, measuring ∼ 1nm in diameter is not large enough to allow the passage of the genome associated protein VPg in its folded state. NMR structure analysis of FCV VPg has shown it to form a compact core of three short α-helices, that measures 2-3 nm in diameter [33]. Further structural changes and possibly the action of an unknown cofactor may therefore be required to trigger injection of the viral genome through the portal vertex and into the cytosol. Such structural rearrangements may require insertion of VP2 into the endosomal membrane or follow environmental cues during endosome maturation. In the course of our investigation of the structure of FCV strain F9, we identified a putative potassium binding site in VP1, found in a region that is important for virus entry. In addition to acidification, endosome maturation is accompanied by considerable ion flux [27], further experiments to explore the role of metal ions in entry and genome release may therefore prove fruitful.

We have applied newly-developed image processing methods to discover and determine the structure of a novel portal assembly in a model calicivirus – FCV [34-36]. Previous attempts to resolve asymmetric features in several icosahedral viruses have proven challenging and largely intractable. This is owing to the weak signal of such features and super-position of density from symmetry related capsomeres in cryoEM projection images. The methods that we have applied in this study are broadly applicable and will allow us to determine for the first time and fairly trivially, the high-resolution structures of biologically important asymmetric components of viruses. The methods employed herein therefore open the door to many more exciting discoveries in virus biology.

Our discovery of a portal assembly at a unique vertex in FCV represents a major step-forward in our understanding of calicivirus biology. We have assigned both structure and function to the poorly understood but indispensable minor capsid protein VP2.

The implications of this discovery extend beyond the *Caliciviridae* however. While endosome escape in enveloped viruses has been well characterised both structurally and biochemically, there are limited data on mechanisms of endosome escape in non-enveloped viruses (reviewed [37, 38]). Some large viruses, such as the DNA containing adenoviruses and double-stranded RNA containing reoviruses require to release their intact nucleocapsid into the cytosol and are thought to induce fragmentation of the endosomal membrane. The small icosahedral positive-sense RNA containing *Picornaviridae* are however thought to release their genomes into the cytosol by the formation of a pore in the endosomal membrane following externalisation of the N-terminus of the capsid protein VP1 and release of VP4 [39]. Our discovery of a large portal assembly in the *Caliciviridae* provides a paradigm for further mechanistic studies of genome translocation across the endosomal membrane that is readily probed by mutational, biochemical and structural analysis. We anticipate that further investigation will yield broadly applicable insights into RNA virus entry mechanisms.

## Materials and Methods

### Virus culture and purification

Feline calicivirus strain F9 was propagated in Crandell Reese Feline Kidney cells (CrFK) for 16 hours. The cell debris was pelleted from the culture medium by centrifugation (1,500xg, 10 min at 4°C). The virus particles within the supernatant were pelleted by centrifugation at 20,000rpm for 2 hours at 4°C using a Surespin630 rotor. The pellets were resuspended in phosphate buffered saline (PBS) and the virus purified by centrifugation through a caesium chloride gradient (1.31-1.45g/ml) using a SW-41 Ti rotor at 28,000rpm for 8 hours and at 12°C. The purified virus particles were extracted from the gradient and pelleted by centrifugation at 20,000rpm for 2 hours at 4°C using an SW-41 Ti rotor. The resultant pellet was then resuspended in 100μl PBS and stored at 4°C until use.

### Expression and purification of fJAM-A

The expression and purification of the soluble fJAM-A ectodomain was performed as previously described[20]. RNA was isolated from CrFK cells and the sequence encoding the signal peptide and ectodomains of fJAM-A was amplified by reverse transcription-PCR (RT-PCR). The PCR product was used to generate a eukaryotic expression plasmid termed pDEF:fJAM-A:Fc which contained the extracellular domains of fJAM-A with the Fc domain of human IgG1 fused at the C terminus. The fJAM-A and Fc domains were separated by a factor Xa cleavage site to allow downstream removal of the Fc tag. Chinese hamster ovary cells stably expressing the soluble fJAM-A:Fc fusion protein were generated and soluble fJAM-A:Fc protein purified from tissue culture supernatant using protein A Sepharose. Monomeric fJAM-A was generated by factor Xa cleavage of fJAM-A:Fc, removal of factor Xa by using Xarrest agarose (Novagen), and removal of the released Fc domain by protein A dynabeads (ThermoFisher).

### Sequencing of FCV VP1 and VP2 ORFs

The sequences for the FCV capsid protein ORFs were determined to facilitate atomic model building. Reverse transcriptase polymerase chain reaction (RT-PCR) was used to isolate the relevant DNA for sequencing using the Sanger method, performed at the MRC Protein Phosphorylation Unit, University of Dundee.

Primers were designed based on published FCV Strain LLK VP1 and VP2 sequence data (Genbank U07131[40]) as follows: VP RT primer: 5’-ttaggcgcaggtgcggcagc-3’ VP1 PCR1 5’-atgtgctcaacctgcgctaacgtgct-3 VP1 PCR2 5’-tcataacttagtcatgggactcctaa-3’ VP2 PCR1 5’-atgaattcaatattaggcctgatt-3’ VP2 PCR2 5’-aattttaaacaaatttttatatga-3’

RNA was extracted from 140 µl of FCV strain F9 infected cell culture medium using Qiagen QiaAMP Viral RNA mini kit according to the manufacturers protocol. The purified RNA was reverse transcribed using Superscript IV RT kit (Thermo Fisher) and VP RT primer in a 20 µl reaction volume according to manufacturer’s protocol. PCR was carried out on 6 µl of the RT reaction using the Q5 High Fidelity PCR kit (NEB). VP1 was amplified using primers VP1 PCR1 and VP1 PCR2, and VP2 was amplified using primers VP2 PCR1 and VP2 PCR2. PCR products were purified using agarose gel electrophoresis and the Geneclean Turbo kit (MPBio).

### Negative stain electron microscopy

FCV particles, both undecorated and labelled with soluble fragments of fJAM-A, were imaged by negative stain transmission electron microscopy. 5 µl of virus preparation was loaded onto a freshly glow-discharged continuous carbon TEM grid. The grid was incubated for two minutes and then washed in three 20µl droplets of water followed by two 20µl droplets of 2% uranyl acetate. Finally the grid was incubated in a 20µl droplet of 2% uranyl acetate for five minutes drained and dried at room temperature. The grids were then imaged in a JEOL 1200 EX II transmission electron microscope equipped with a GATAN Orius camera.

### Cryo-electron microscopy

Purified FCV particles were incubated with the soluble ectodomain of fJAM-A for 1 hr at 4°C. The undecorated and decorated virions were then prepared for cryo-electron microscopy by loading 5µl onto freshly glow-discharged, carbon coated c-flat holey carbon grids (CF-22-4C, Protochips Inc.) in a Vitrobot vitrification robot (FEI) held at 4°C and 100% humidity. Grids were blotted for 4 seconds prior to being frozen by plunging into a bath of liquid nitrogen-cooled liquid ethane.

Vitrified specimens were imaged at low temperature in a Thermo-Fisher Titan Krios equipped with a Falcon III detector. The column was operated at 300 keV accelerating voltage and a nominal magnification of 75,000x resulting in a pixel size of 1.065 Å/pixel. Each micrograph was recorded as a movie of 50 individual fractions with a total dose of 63 e/Å^2^. Automated data-collection was performed using EPU software. Data collection was performed at the Astbury BioStructure Laboratory, University of Leeds.

### Three-dimensional image reconstruction

Micrograph stack files were corrected for drift using MotionCor2[41] and defocus estimation performed by Gctf[42] as implemented in Relion 2.1[43]. 59,531 undecorated particles were automatically picked from 5198 micrographs and 129,884 fJAM-A decorated particles from 13,865 micrographs. Particle images were subjected to both 2D and 3D classification to exclude false-positives and damaged particles. Final datasets of 41,436 undecorated and 71,671 fJAM-A decorated virions were subjected to 3D refinement to calculate final icosahedrally averaged three-dimensional reconstructions. These were the subjected to post-processing and local resolution analysis in Relion 2.1 to determine both final resolution and optimal sharpening parameters. Maps were visualised using UCSF Chimera [25] and ChimeraX and coloured according to local resolution, unless otherwise specified.

### Model based classification

To attempt to separate fJAM-A decorated FCV virions into separate classes based on the extent of conformational change they had undergone, unsupervised 3D classification was performed in Relion 2.1 using our previously published structures as starting models[21].

### Focused classification

To resolve the structures of individual asymmetric features we employed a focused classification approach in Relion 2.1 [34-36]. Briefly, following 3D refinement with imposition of icosahedral symmetry, the symmetry of the data set was expanded such that each particle was assigned 60 orientations corresponding to its icosahedrally redundant views. A cylindrical mask was prepared to exclude all of the capsid structure except the area of interest, using SPIDER [44]. Unsupervised 3D classification was then performed without orientation refinement. Thus, a single capsomere or symmetry axis was reconstructed, and classification was performed to resolve structural differences.

### Atomic model building

Atomic models were built using Coot [45], the CCP-EM [46] suite and Phenix [47]. Briefly, the structure for VP1 of FCV in the undecorated virus was built starting from our previously calculated homology model [20]. The structure was manually edited in Coot to correct the sequence and improve the correspondence to our EM density map. This was followed by rounds of iterative real-space refinement in Phenix. To build an atomic model of VP1 in our receptor-bound models, the P-domains and S-domains were docked independently as rigid bodies using UCSF Chimera [25]. The models were then manually edited and iteratively refined as above. Likewise, the structure for fJAM-A was built starting from our previously calculated homology model and refined using Coot and Phenix. The structure of VP2 was manually built ab Initio in Coot and refined using Phenix. Model validation was performed using MolProbity [48], as implemented in Phenix. Modelling of metal ions was validated using the checkmymetal server (https://csgid.org/csgid/metal_sites) [24].

### RNA release assay

Equal concentrations of FCV and fJAM-A ectodomain were combined in Tris buffer of the appropriate pH. 10µl Syto9 nucleic acid binding dye (ThermoFisher Scientific) was added to a final volume of 100µl in a black 96-well plate. For positive controls, 550 ng FCV RNA was added in place of the FCV and fJAM-A samples. Fluoresence readings were collected every 5 minutes over a period of 4 hours using a PHERAstar FS (BMG labtech) plate reader equipped with a 485/520 filter cube. Data were normalised to the corresponding samples containing only FCV and Syto9.

### Small Angle X-ray Scattering (SAXS)

The fJAM-A ectodomain was dialysed into PBS containing 0.01% sodium azide, 1% sucrose and 10 mM potassium nitrate and samples taken to the Diamond Light Source (Oxfordshire, UK) where high-pressure liquid chromatography (HPLC) small angle X-ray scattering (SAXS) data were acquired. The HPLC column (Superdex 200 (GE Healthcare)) was equilibrated with buffer for 1 hour, the sample loaded onto the column from a 96-well plate and the diffraction data collected. Eight 60-second frames of data, for which the radius of gyration (Rg) was observed to be constant, were selected for further processing with PRIMUS [49]. A Guinier region was observed between 0.26 ≤ qRg ≤ 1.29 Å2 (where q is the momentum transfer (4πsin(Φ)/λ)) giving Rg = 30.7 Å. The particle distance distribution function (p(r) versus r, where r is the real-space distance) was evaluated using GNOM [50] with a maximum dimension of 102.8 Å. An ab initio envelope was computed from p(r) vs r using DAMMIF [51]. The dimeric model of human fJAM-A ectodomain (PDB: 1NBQ) was fitted into the SAXS envelope using UCSF Chimera [25].

### Data deposition

The icosahedral reconstruction of undecorated FCV and the C3-symmetrised reconstruction of FCV-fJAM-A are deposited in the EM databank with accession numbers EMD-0054 and EMD-0056 respectively. The atomic coordinates for the FCV capsid asymmetric unit (VP1) are deposited in the protein data bank with accession number PDB-6GSH. The atomic coordinates for the FCV-fJAM-A portal vertex (VP1, VP2 and fJAM-A) are deposited in the protein data bank with accession number PDB-6GSI. Motion corrected micrographs of undecorated and fJAM-A labelled FCV (the raw data) are deposited in the EMPIAR data bank with accession numbers EMPIAR-10192 and EMPIAR-10193 respectively.

## Acknowledgments

CryoEM data in this study were collected at the University of Leeds, Astbury BioStructure Laboratory. The authors thank and acknowledge Sjors Scheres for advice on the application of focussed classification in Relion, Rebecca Thompson for her expert microscopy support, Neil Ranson for microscope access and discussions, Joseph Hughes for advice on statistical analysis, Peter Stockley, Massimo Palmarini and John McLauchlan for their input during the preparation of this manuscript. We acknowledge Diamond Light Source for time on Beamline B21 under Proposal MX11651-24. M.J.C. was supported by a PhD studentship from the UK Biotechnology and Biological Sciences Research Council (BBSRC WestBIO DTP: BB/J013854/1). D.B and M.M. are supported by the UK Medical Research Council (MC_UU_12014/7).

